# Bacteriophage therapy for the treatment of *Mycobacterium tuberculosis* infections in humanized mice

**DOI:** 10.1101/2023.01.23.525188

**Authors:** Fan Yang, Alireza Labani-Motlagh, Josimar Dornelas Moreira, Danish Ansari, Jose Alejandro Bohorquez, Sahil Patel, Fabrizio Spagnolo, Jon Florence, Abhinav Vankayalapati, Ramakrishna Vankayalapati, John J. Dennehy, Buka Samten, Guohua Yi

**Author notes:** Corresponding authors. Guohua Yi, Buka Samten, and John J. Dennehy. Contribute equally (Fan Yang, Alireza Labani-Motlagh).

## Abstract

The continuing emergence of new strains of antibiotic-resistant bacteria has renewed interest in phage therapy; however, there has been limited progress in applying phage therapy to multi-drug resistant *Mycobacterium tuberculosis* (*Mtb*) infections. In this study, we tested three bacteriophage strains for their *Mtb*-killing activities and found that two of them efficiently lysed *Mtb* H37Rv in 7H10 agar plates. However, only phage DS6A efficiently killed H37Rv in liquid culture and in *Mtb*-infected human primary macrophages. In subsequent experiments, we infected humanized mice with aerosolized H37Rv, then treated these mice with DS6A intravenously to test its *in vivo* efficacy. We found that DS6A treated mice showed increased body weight and improved pulmonary function relative to control mice. Furthermore, DS6A reduced *Mtb* load in mouse organs with greater efficacy in the spleen. These results demonstrated the feasibility of developing phage therapy as an effective therapeutic against *Mtb* infection.

---

In 2020, while all focus was on COVID-19, *Mycobacterium tuberculosis* (*Mtb*), the causative pathogen of Tuberculosis (TB), infected 10 million people and caused 1.5 million deaths globally [1], making it one of the leading causes of death by a single infectious agent. More importantly, one-third (>2 billion) of the world’s population has been latently infected with *Mtb* (LTBI) [2]. When complicated with other diseases/infections, such as HIV and diabetes, these co-morbidities can cause LTBI patients to develop active TB and increase mortality [3]. Although these devastating facts highlight the threat posed by this deadly bacterium, the emergence of drug resistant *Mtb* strains in recent years has worsened the situation in terms of *Mtb* prevention and control.

Upon establishment of infection, *Mtb* exhibits remarkable abilities to adapt to the local environment to evade the host’s immune responses due to its unique and dynamic four-layer cell envelope structure, which can help it adapt to hostile lung microenvironments and facilitate its entry into a non-replicating drug-tolerant persister state. In this state, the bacilli are well protected against antibiotic therapy [4, 5], and are potentially a source of new strains of drug-resistant *Mtb* (DR-TB)[6]. In 2019, around half a million people developed DR-TB, and 78% of these cases developed multidrug-resistant TB (MDR-TB) or extensively drug-resistant TB (XDR-TB), which are resistant to the first-line drugs, rifampicin, and isoniazid, or to second-line drugs [1]. MDR-TB and XDR-TB are more challenging to treat and are associated with increased morbidity and mortality [7]. Therefore, the development of alternative therapies in addition to antibiotics is critical for the advancement of TB therapy in the era of *Mtb* drug resistance.

Phage therapy has emerged as a renewed approach to eliminating bacterial infections [8, 9]. Phages, viruses that infect bacteria, are bacteria’s natural enemies and have been used to control bacterial infections since even before the discovery of antibiotics [10]. In the age of antibiotic resistance, phage therapy has drawn tremendous attention. Recent clinical cases and trials demonstrated that phages can be used to treat antibiotic-resistant bacterial infections with positive clinical outcomes [11–16], and patients with drug-resistant *M. abscessus* and *M. chelonae* have also been successfully treated [17–19]. These prominent bodies of work on using phage therapy to treat other mycobacterial infections inspired us to pursue phage therapy as a potentially effective treatment for *Mtb* infection. However, as to the phage therapy for *Mtb* treatment, the significant advancements have focused on screening lytic phage strains for effective killing of *Mtb* in bacterial culture plates, demonstrated by plaque formation [20–22], but no systematic study has characterized the effectiveness of phages in killing *Mtb* in primary human macrophages or in reliable animal models. Therefore, extensive preclinical studies, especially using primary human macrophages or animal models that resemble human clinical settings, are of exceptional clinical importance in the development of phage therapy for *Mtb* treatment.

One of the major challenges to eliminating *Mtb* infection is *Mtb*’s induction of granuloma formation once host defenses fail to kill the bacteria. While granuloma formation limits *Mtb* growth, it provides a survival niche for *Mtb* replication when the immune system is weakened. Granulomas, the hallmark of TB pathology, consist of macrophages, neutrophils, and lymphoid cells, including T and B cells, and are formed upon *Mtb* infection in patients [23]. The commonly employed mouse model can be infected with *Mtb*, but *Mtb* can only form a granuloma-like structure in mice. Mouse *Mtb* infections lack the caseous necrotic granulomas that are often observed in TB patients [24], suggesting human immune cells may be needed to recapitulate real clinical settings. Recently humanized mice based on NOD-scid IL2Rgamma^KO^ (NSG) mice were developed. However, the mouse innate immune system in these mice was deficient and transplanted human hematopoietic stem cells (hHSCs) were generally not well-developed; thus, the human B, T, and myeloid cells were immature, and NK cells lost functions [25, 26]. These mouse models provide valuable tools to study *Mtb* infection [24, 27]; however, the myeloid barrier in these NSG mice causes a relative lack of leukocyte differentiation [26]. The difficulty of forming sufficient granulomas (which need mature macrophages) makes these mice less able to recapitulate *Mtb* infection in humans.

Nevertheless, the newly developed NSG-SGM3 mice transgenically express three human cytokine/chemokine genes IL-3, GM-CSF, and KITLG, which can enhance the differentiation and maturation of the myeloid-lineage cells [26, 28–31]. Moreover, these three transgenic genes can improve the homeostasis of human CD34+ HSCs, increase neutrophil and macrophage numbers and function, and stimulate hCD34+ cells to differentiate into myeloid progenitor cells [30, 32]. Thus, this humanized mouse model can generate sufficient and fully functional myeloid cells, including macrophages, to study *Mtb* infection in the context of an entire human immune system.

In this study, we show that phage DS6A can efficiently kill wild-type *Mtb* H37Rv *in vitro* (in both 7H10 agar plates and 7H9 liquid culture), in primary human macrophages, and in humanized NSG-SGM3 mice, demonstrating the feasibility of developing phage DS6A as an effective therapeutic against *Mtb* infection.

## Results

### Phage infectivity in *M. smegmatis* and in *M. tuberculosis* H37Rv

We first tested three bacteriophages, D29, Chah and DS6A, for their lysis capacities of *M. smegmatis* and *Mtb* H37Rv by plaque assay. Serially diluted phages (10^2^-10^4^ pfu) were incubated with *M. smegmatis* or H37Rv in 12-well plates with 7H10 agar. The results showed that two phage strains, D29 and Chah lysed *M. smegmatis* (Fig. 1a) and formed different sizes of plaques, and two phage strains D29 and DS6A, lysed H37Rv and formed plaques (Fig. 1b).

**Figure 1.**
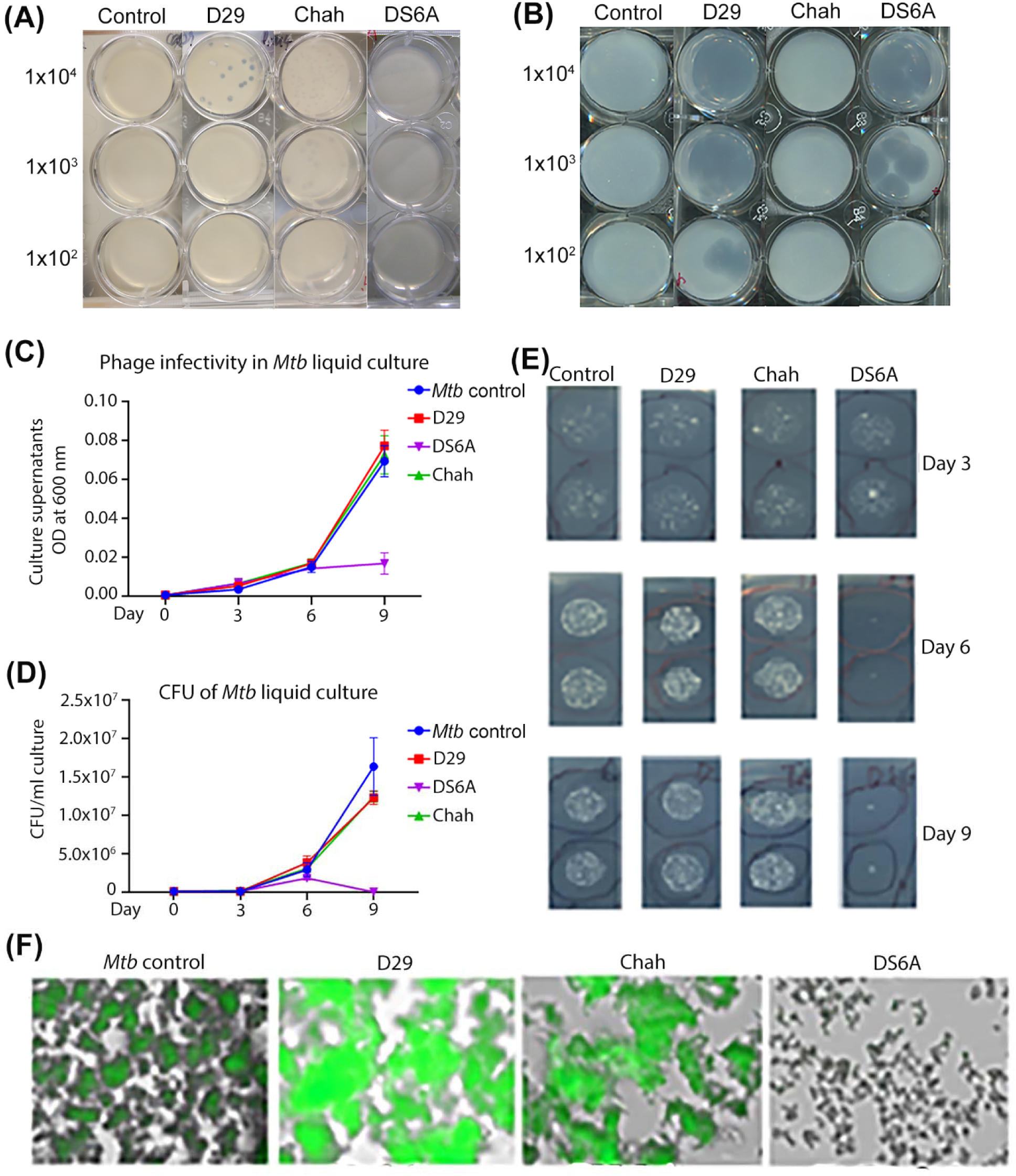
Phage infectivity of *M. smegmatis* and *Mtb* H37Rv in solid agar plates, and in *Mtb* liquid culture. **(a) and (b)**: Serial dilutions (10^2^-10^4^ pfu) of different strains of bacteriophages were mixed with 1 x 10^5^ CFUs *M. smegmatis* **(a)** or H37Rv **(b)** in 7H9 medium and incubated at 37°C with shaking for 1 h. The infection cultures were then mixed with 0.8% top agar and spread on 7H10 agar plates supplemented with 10% OADC enrichment. **(c to f)**: H37Rv (1 x 10^5^ CFUs) were infected with various bacteriophages at an MOI of 1 for 1 h, then inoculated into 20 mL 7H9 media supplemented with 10% ADC and incubated at 37°C for 9 days. The cultures were sampled at days 3, 6, and 9 post infection by plating 10 μL of 10-fold serial dilutions of on the 7H10 agar plates and cultured at 37°C for CFU determination. **(c):** OD_600_ of sampled liquid cultures was measured at different time points as indicated. **(d):** Statistics of the CFUs of each sampled liquid culture sampled at various time points. **(e):** Titering assay of phage-*Mtb* liquid culture sampled at different time points. The data are representative results of two independent experiments. **(f):** Phage infection of H37Rv that expresses a GFP reporter gene. An MOI of 1 was used to infect H37Rv-GFP. GFP expression was pictured under fluorescence microscopy after nine days of phage infection.

We then tested the *Mtb* killing ability of these three phage strains in liquid culture by measuring the optical density at 600 nm (OD_600_) of the H37Rv suspensions at different time points after incubation with phages at one multiplicity of infection (MOI, ratio of phage to bacteria) by inoculation of 7H9 media with 5×10^4^ CFUs of H37Rv and 5×10^4^ pfu of phages. We found that only DS6A significantly reduced *Mtb* OD_600_ on day 9 (Fig. 1c), while other phages did not affect *Mtb* OD_600_ on day 9, although D29 also showed the ability to kill *Mtb* in agar plates. We then plated the cultures from different time points to determine the CFU of the bacilli (Fig. 1d and 1e). The CFU results were consistent with the OD_600_ measurements. The above results also indicated that the *Mtb* killing ability in agar plates does not necessarily correlate with the *Mtb* killing ability in liquid culture.

We further confirmed this outcome using a GFP-expressing H37Rv strain, H37Rv-GFP, in a liquid culture condition as described above. Due to the highly efficient expression of GFP gene, the live bacteria are visible under fluorescent microscopy. The microscopy images of the day 9 cultures clearly showed that there was no GFP expression in the wells with H37Rv-GFP co-cultured with DS6A, whereas H37Rv-GFP in the wells co-cultured with other phages remain fluorescent, further demonstrating the highly effective *Mtb*-killing ability of DS6A (Fig. 1f).

### Phage DS6A can efficiently eliminate *Mtb* H37Rv in primary human macrophages

To test whether the phages can kill *Mtb* in infected primary human macrophages, which is critical for therapeutic effectiveness in clinical settings, we isolated peripheral blood mononuclear cells (PBMCs) from the blood of a healthy donor and isolated the hCD14+ monocytes using microbeads to above 95% purity as determined by flow cytometry analysis (Supplementary Fig. 1) and differentiated the purified CD14+ cells into macrophages by incubation of the cells with hM-CSF, hGM-CSF, and hIL-4 for 5 days. The macrophages were then infected with H37Rv and treated with different phages. On days 5 and 10, we plated the macrophage lysates on 7H10 agar plates to determine the CFUs (Fig. 2a). The results showed no significant reduction in *Mtb* CFUs for all phage-treated groups on day 5 when compared to the *Mtb*-infected macrophages without phage treatment as a control. However, on day 10, phage DS6A completely eradicated the bacilli from the infected macrophages, while other phages still failed to show any reduction in *Mtb* growth in macrophages (Fig. 2b and 2c).

**Figure 2.**
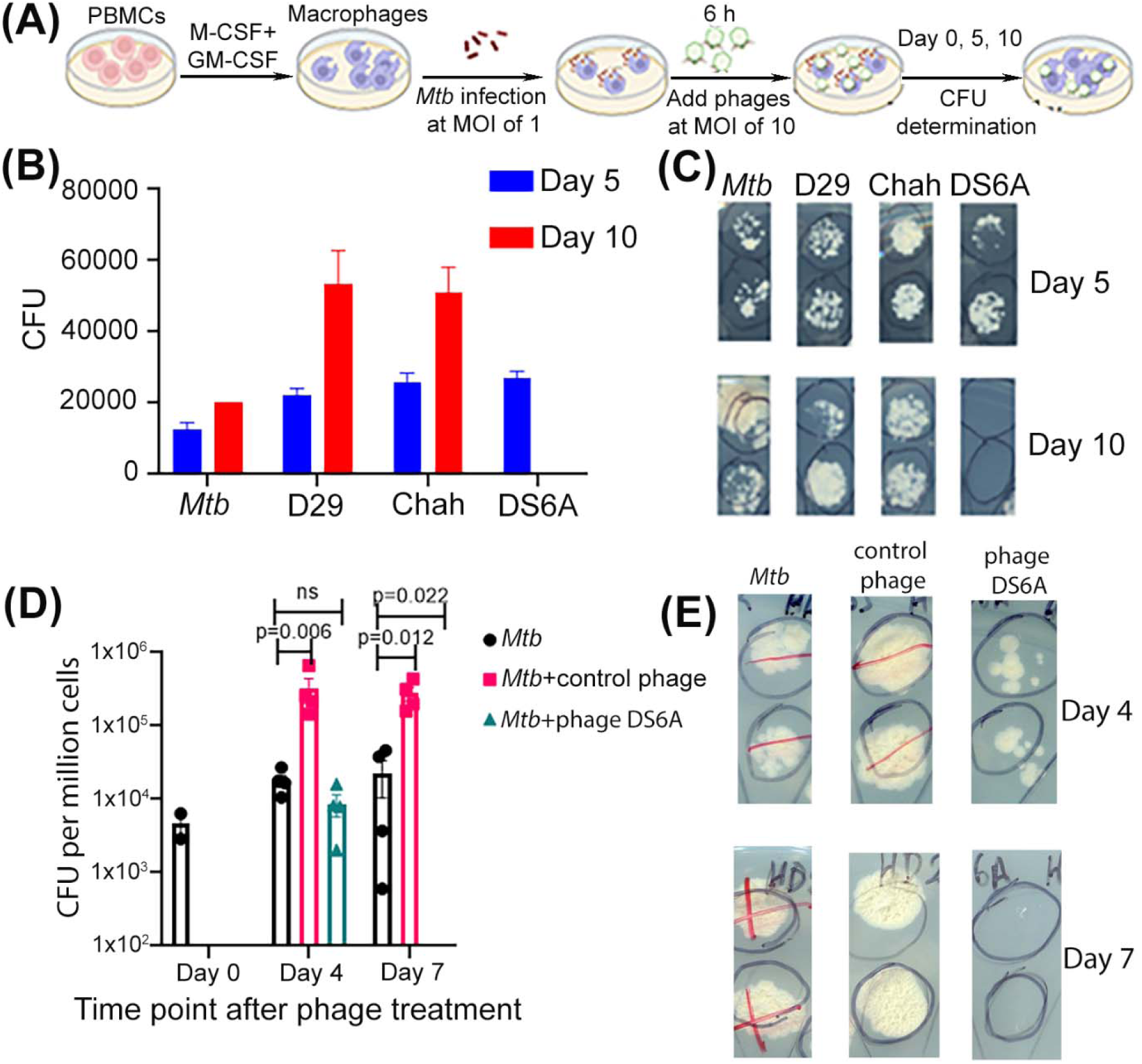
Phage DS6A eliminates *Mtb* in infected primary human macrophages. **(a)** Experimental procedures of the assay. Panels **(b)** and **(c):** Half million microphages were infected with H37Rv at an MOI of 1. After 4 h of *Mtb* infection, three different phages (5×10^6^, MOI of 10) were applied to the *Mtb*-infected primary human macrophages, and cultured at 37°C. The cells were sampled at days 0, 5 and 10 to determine the bacillary load for the evaluation the *Mtb*-killing ability of the phages by plating the ultrasound-broken cells on 7H10 plates (supplemented with 10% OADC) and the CFUs were counted on Day 8. Panel **(b)** shows the statistics of the CFUs of all phages, and panel **(c)** shows the representative pictures (duplicated) of the 7H10 plates taken at day 5 and day 10. Panels **(d)** and **(e):** Testing the *Mtb*-killing ability of phage DS6A in *Mtb*-infected macrophages derived from four different healthy donors, and phage D29 was used as a negative control. The infected macrophages were sampled at different time points, sonicated, and plated on 7H10 agar plates using different dilutions to titer the bacillary load. Panel **(d)** shows the statistics of five donors, and panel **(e)** shows the representative pictures taken from 10x diluted plates for days 4 and 7 cultures.

We further tested whether the phages can also kill *Mtb* in infected macrophages from different donors. For this purpose, we isolated monocytes from four healthy donors’ PBMCs, and repeated the above experiment in these four samples using phage DS6A, and phage D29 was used as a negative control. Meanwhile, we wanted to see whether *Mtb* could be killed in a shorter time rather than ten days, so we set the sampling time points to days 4 and 7. The results showed that all four donors displayed the same pattern in which *Mtb* in the infected macrophages were killed within seven days by phage DS6A, but not in the control phage treatment (Fig. 2d and 2e). Interestingly, some of the phage-treated cultures have a significantly higher bacillary load than the control bacteria cultures (Fig. 2b and 2d), which we address further in the discussion section.

### *Mtb* infection of humanized NSG-SGM3 mice

Compared to the commonly used Hu-PBL and BLT mouse models, the humanized NSG-SGM3 mouse model has several advantages: 1) Humanized NSG-SGM3 mice exhibit improved reconstitution with an increased general population of human immune cells (hCD45+); 2) Cytokine expression supports human myeloid cell differentiation and maturation; thus the functional macrophages and dendritic cells are significantly elevated in the bone marrow compared with NSG recipients [26, 29]; 3) There is a significant increase of regulatory T cells (Treg) [29], which are important in regulating *Mtb* pathogenesis [33, 34]; 4) B cells can be developed to mature phenotypes with the ability of class-switching [25], which is important to evaluate the IgM and IgG responses against phages; 5) They are relatively easy to generate by intravenous injection of human HSC into irradiated adult mice for humanization.

We obtained NSG-SGM3 mice from the Jackson Laboratory and are breeding them in-house. For humanization, 4-6-week-old NSG-SGM3 mice were irradiated at 150cGy and then infused with human CD34+ HSC. Ten to fifteen weeks after HSC injection, the animals developed a full complement of human immune cell types, including CD4+ and CD8+ T cells, B cells, myeloid cells, and natural killer (NK) cells (Fig. 3a).

**Figure 3.**
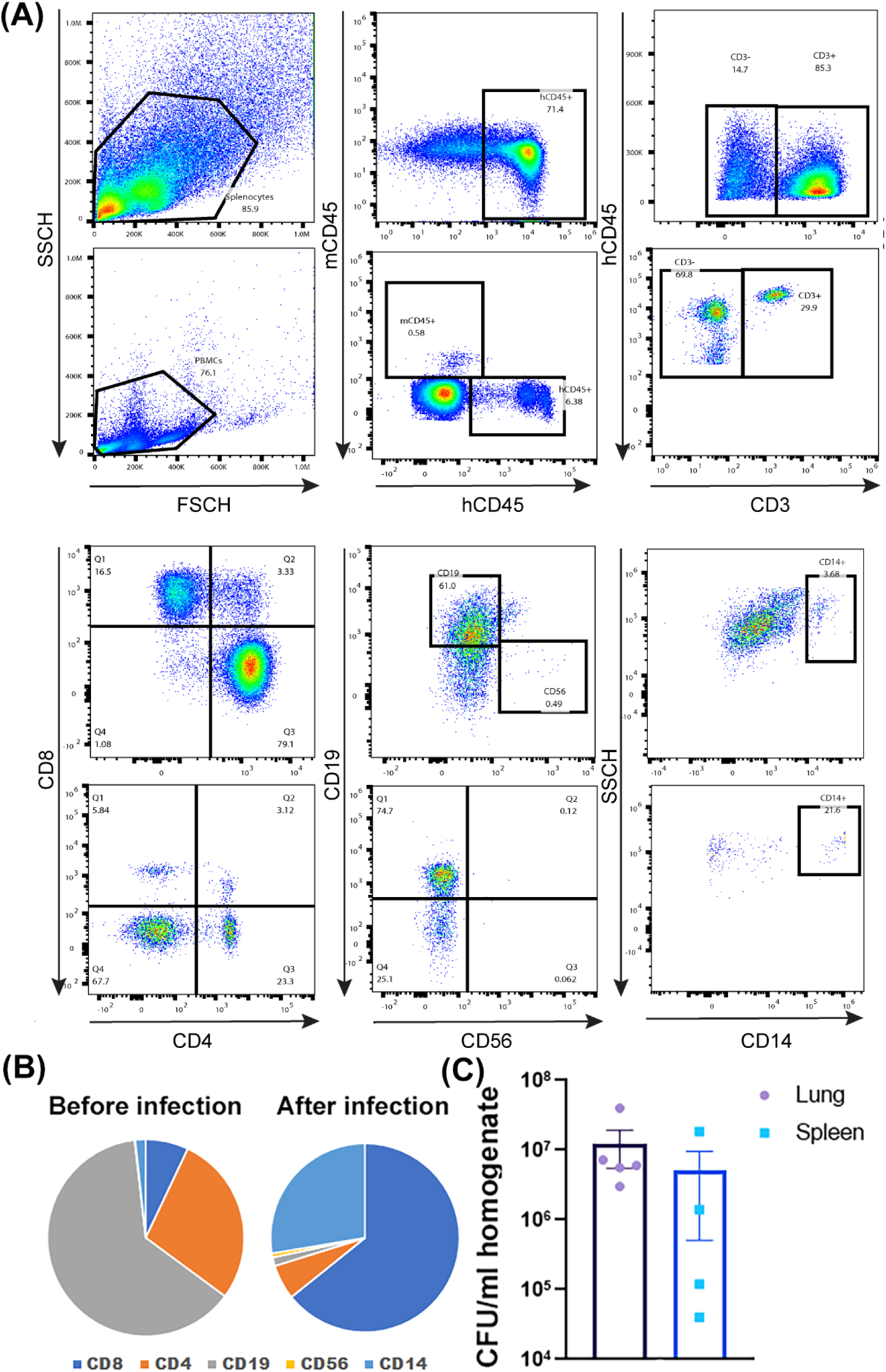
*Mtb*-infection of humanized NSG-SGM3 mice. **(a)** Reconstitution of human immune cells in humanized NSG-SGM3 mice. NSG-SGM3 mice were irradiated and intravenously injected with 2 × 10^5^ hCD34+ HSC. After 10-12 weeks, reconstitution of human T cells, B cells, myeloid cells, and natural killer (NK) cells within human CD45-gated population was analyzed by flow cytometry. **(b)** Statistics of different immune cells before and after infection. **(c)** Five mice were sacrificed to check *Mtb* infection in lungs and spleens by determining the organ CFUs by plating serially diluted organ homogenates on 7H10 agar plates as described above.

To establish humanized NSG-SGM3 mouse model of *Mtb* infection, we infected the humanized NSG-SGM3 mice with H37Rv via a Madison chamber to deposit about 50-100 CFUs per mouse lung. At 25 days post-infection, we sacrificed the mice and used the homogenates from the lungs and the spleens to analyze the immune cell profile after infection. Additionally, we determined *Mtb* growth in lungs and spleens. The results showed that, when compared with the PBMCs analysis before infection, the spleen immune cells have undergone substantial changes after infection, and the CD8+ T cells and myeloid cells have significantly expanded (Fig. 3b). Meanwhile, from both the lung and spleen homogenates, we can detect a high number of bacilli (Fig. 3c), indicating that the mice established the *Mtb* reservoirs not only in the lungs but also systematically.

### Phage DS6A effectively eradicates *Mtb* in humanized mice

To investigate if phage DS6A can eradicate *Mtb in vivo*, we performed the animal experiment in the humanized NSG-SGM3 mouse model of tuberculosis (Fig. 4a). We infected the humanized NSG-SGM3 mice with low dose aerosolized H37Rv and as confirmed by an average ∼100 bacilli per lung deposited at day one post infection (Fig. 4c). From day 3 of *Mtb* infection, we treated the mice every other day with ten doses of DS6A (10^11^ phage particles/dose) via intravenous administration.

**Figure 4.**
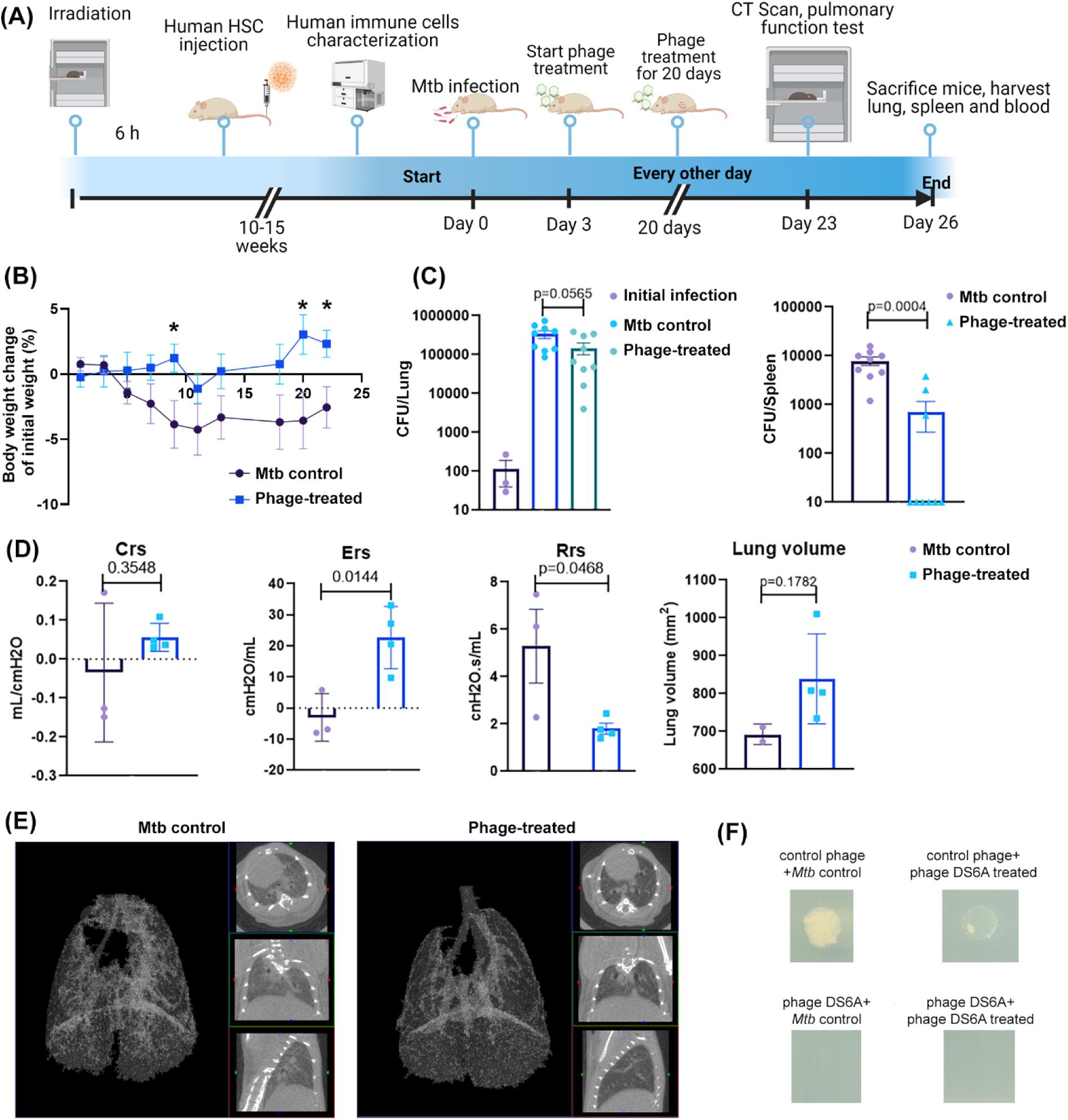
Phage DS6A is effective at eradicating *Mtb* in humanized NSG-SGM3 mice. **(a)** Schematic presentation of the animal experiment procedures. H37Rv aerosol infected humanized mice were treated three days after infection with 1×10^11^ pfu of phage DS6A per mouse or equal volume of sterile PBS every other day for eight times. **(b)** The mouse body weights were monitored over time, and the body weight change over the initial body weight of the mice are shown as percentages**. (c)** The growth of H37Rv in the mice lung and spleen homogenates together with initial infection dose (left panel in purple) are shown. **(d)** Before termination of the experiment, the mice were subjected to pulmonary function testing. The measurements of elastance, compliance, total lung resistance, and total lung volume were collected, and the statistics are shown. **(e)** CT scans were performed for each mouse, and two representative figures show the *Mtb*-infected control mouse and phage-treated mouse lungs, respectively. The left panel of each mouse CT scan figure shows the 3D image, the white areas represent the high-density scan (e.g., tissues), while the black areas represent low density scan (e.g., air). The three small figures in each mouse scan show different angles of scan results. **(f)** The colonies from the plates that spread with lung or spleen homogenates of the H37Rv infected control or phage-treated mice were picked and mixed with 1 x 10^6^ pfu of phage DS6A or a negative control phage (phage Chah) in 10 µl 7H9 media and incubated at 37°C for 1h. Then the infection cultures were pipetted onto the 7H10 agar plates OADC and incubated at 37°C for determination of CFUs. The animal experiments were performed four times, and the 3^rd^ and 4^th^ experiments were shown. Except the pulmonary functions were tested once (3^rd^ animal experiment), the other data are shown as the combined results from the 3^rd^ and 4^th^ experiments. All the statistics are shown as mean ± SEM.

After the treatments, we determined the growth of *Mtb* in mouse lungs and spleens, performed pulmonary function tests (PFTs) and overall health evaluation by monitoring mouse body weight changes in association with changes in bacillary burden of the mice. The results showed that DS6A-treated *Mtb*-infected mice gained significantly more weight than control *Mtb*-infected mice (Fig. 4b). The PFTs also showed that two of the four parameters tested in the phage-treated group had significantly improved, with less respiratory resistance and better elasticity, demonstrating improved pulmonary function after treatments (Fig. 4d). The CT scan results showed that there were fewer high-density areas with much cleaner lungs in the phage-treated group of mice, which indicates less inflammation and other pathological changes (Fig. 4e). Furthermore, the lung volume of the phage-treated mice was larger than untreated *Mtb* control mice, even though the difference didn’t reach significance (p=0.1782) (Fig. 4d). The CFU counts showed that the bacilli were completely eradicated in the spleens in six of nine mice of the phage-treated group, and the average bacillary load in the spleen was significantly lower than in the *Mtb*-infected control group (Fig. 4c). Moreover, the lung bacillary load showed three-times higher in *Mtb*-infected control mice than in phage-treated mice, while the difference didn’t reach significance (Fig. 4c). These results suggest that, after intravenous administration, phage DS6A has the capacity to kill the replicating *Mtb* within/or outside the lung that are disseminated to the blood circulation.

To explain why the phage DS6A performed better in killing *Mtb* bacilli in spleens than in lungs, we determined the phage genome copies in the lung and spleen homogenates. The quantitative PCR result showed that the spleen had three times more phage copies in spleens than in lungs (Supplementary Fig. 2), suggesting that the effectiveness in tissues may be positively correlated to the phage copies, and that phages intravenously administered preferentially distributed and replicated more in the spleen than in the lungs, which may explain the organ-dependent differences in *Mtb* eradication in humanized mice.

Given that phage DS6A could not wholly eradicate the spleen bacteria in several mice, we wondered whether phage resistance developed in the bacteria within these mice. For this purpose, we picked colonies from the plates that spread the lung and spleen homogenates of phage-treated mice, and then mixed them with phage DS6A to see if the phage could kill the bacteria *in vitro*, and the phage Chah was used as a negative control. The results showed that the negative control phage could not kill both bacteria colonies picked from DS6A-treated or untreated mice, while the DS6A could kill both (Fig. 4f), indicating that there was no phage resistance developed during the 20-day phage treatment period.

### Phage therapy elicits antibody responses in humanized NSG-SGM3 mice without affecting the function of immune cells

There is always a concern that the immune responses, especially antibody responses, will stymie the effect of phage therapy. Therefore, we determined the phage DS6A-specific antibody responses in sera from our experimental mice by ELISA. We detected a significantly higher level of human IgM in phage DS6A-treated mouse sera (Fig. 5b). However, we could only detect weak IgA and IgG responses in these mice (Fig. 5a and c).

**Figure 5.**
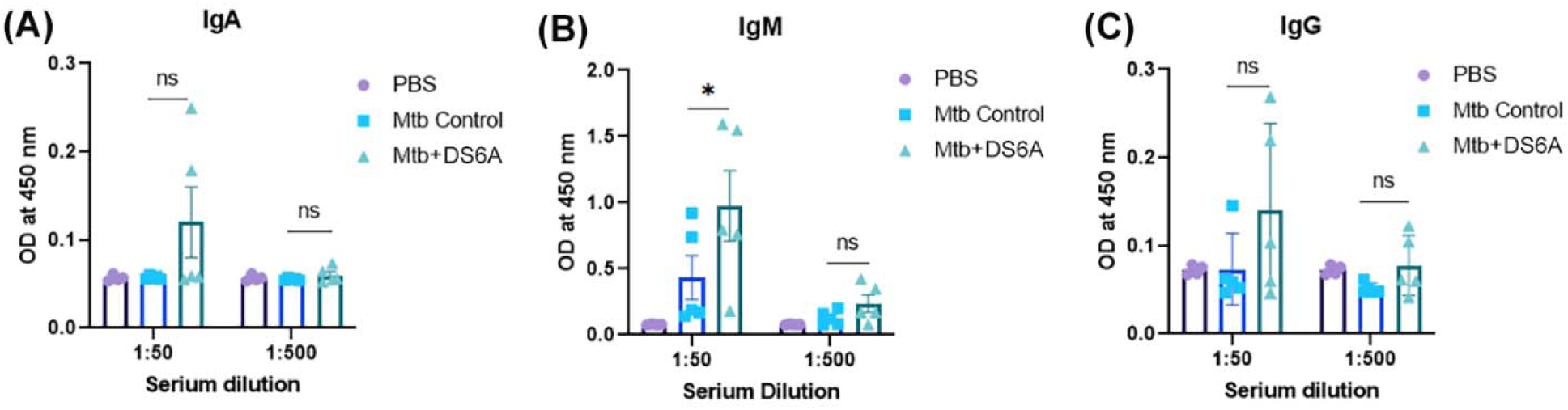
Humanized NSG-SGM3 mice developed antibody responses against phage DS6A. ELISA was performed to determine the phage DS6A-specific human IgA, IgM, and IgG titers in sera of the humanized mice infected with H37Rv and treated with phage DS6A or PBS buffer control. The experiment was repeated once, and the results from one experiment with five mice each group are shown.

We further tested whether phage therapy could affect the cytokine profiles that will be in association with reduced *Mtb* growth and improved lung function due to the phage treatment. We determined the levels of 27 cytokines including cytokines of human lymphocytes and myeloid cells in the lung and spleen homogenates of the mice by a multiplex cytokine assay (Supplementary Fig. 3). Overall, the cytokine results showed that the cytokine levels in the spleen were higher than that of the lungs, and that the phage treatment induced a few myeloid cells-derived cytokines significantly lower than that of the control *Mtb*-infected mice, consistent with reduced bacilli burden in the mice treated with phage. We did not see significantly elevated levels of T cell cytokines in the mice after four weeks of *Mtb* infection, consistent with significantly increased CD14+ monocytes, but not lymphocytes, in the humanized mice after *Mtb* infection.

## Discussion

The development of alternative treatments to antibiotics for *Mtb* infection is an urgent task in the era of antibiotic resistance. In this study, we have shown that bacteriophages D29 and DS6A can efficiently kill *Mtb* bacilli on 7H10 agar plates, while only phage DS6A significantly killed *Mtb* in liquid cultures. Similarly, only phage DS6A was able to kill *Mtb* bacilli in primary human macrophages. In addition, we showed that phage DS6A could effectively eradicate *Mtb* bacilli in spleens and reduced the bacilli load in the lungs in a humanized mouse model of *Mtb* infection. Thus, phage DS6A appears to have the potential to be developed as an effective therapeutic for *Mtb* treatment.

Phage therapy has shown great potential in controlling some intransigent drug-resistant bacterial infections. Compared to typical antibiotic treatment, phage therapy has advantages such as: host specificity, low toxicity, the ability to lyse the bacteria, increased effectiveness (since they can self-replicate *in situ*), and a narrower potential for inducing resistance, all of which render it an attractive treatment strategy in the era of antibiotic resistance. Several phage strains have shown potential in eliminating *Mtb* infection [20]. Unfortunately, there has been little progress in *Mtb* phage therapy, with only two studies published to date that reported on the use of phage therapy for the treatment of *Mtb* in guinea pigs, with limited success [35, 36]. Therefore, our study showing the effectiveness of phage therapy in treating *Mtb* infections of clinically relevant humanized mice represents a significant advancement in this field.

Humanized mouse models provide versatile tools to study various infectious diseases. For *Mtb* infection, the myeloid cell lineage is especially important because the macrophages are the main targets of the bacilli. In most NSG-based humanized mice, myeloid cell differentiation is not efficient, and the macrophages are not mature, thus the physiological processes such as phagocytosis and endocytosis, which may be involved in *Mtb* infection and bacteriophage uptake, may not be completed as properly as they are in humans. However, due to the transgenic expression of the three human cytokine genes, the myeloid cells were well-differentiated in NSG-SGM3 humanized mice that we have used in this study. After *Mtb* infection, the myeloid cells were successfully propagated (Fig. 3b). Moreover, after phage treatments, the phage was shown capable of killing *Mtb* in human macrophages engrafted in the humanized mice, suggesting that NSG-SGM3 mice can recapitulate human *Mtb* infections, and thus NSG-SGM3 mice are an appropriate and clinically relevant animal model for *Mtb* phage therapy.

Although D29 and DS6A showed the ability to kill *Mtb* on bacterial lawns, only DS6A efficiently killed *Mtb* in liquid culture, in human primary macrophages, and in humanized mice, suggesting that DS6A is the most appropriate phage therapeutic for *Mtb* infections. The reasons why D29 can kill H37Rv efficiently on bacterial lawns, but not in liquid culture, are not fully clear (Fig. 1). The most likely explanation is that *Mtb* readily acquires resistance to D29, but not to DS6A. When inoculated in liquid culture at an MOI = 1, it may be that at least some *Mtb* cells are genetically incapable of being infected by D29, thus these resistant *Mtb* cells eventually dominate the culture. Additionally, it appears that D29 cannot easily break this resistance. Nevertheless, these results indicate that liquid cultures, rather than bacteria lawns, appears to be a better prediction model for the *in vivo* efficacy of *Mtb* phage therapy.

Of note, we also observed that once the phage cannot kill *Mtb* in liquid culturing condition, it sometimes will stimulate *Mtb* growth, as shown in Fig. 2. The observation that, in some cultures, bacterial load is higher in the presence of phages D29 and Chah but not DS6A, than in phage-free controls (Fig. 2b and 2d) is puzzling.

Nevertheless, our experiments provide clear evidence that phage DS6A was effective in killing *Mtb* in the spleen via the intravenous route. However, the bacterial clearance in the lungs was not as effective as in spleens. This discrepancy is possibly due to the administration route, as this is one of the most important factors that affects *in vivo* efficacy of phage therapy [37]. Through the intravenous route, phages can easily reach the spleen and liver through circulation as blood filtering organs, but the lungs are much more difficult to reach. A previous study showed that the intravenous delivery of a pneumonia bacteria-specific phage led to a 2-log lower prevalence (<100 times) in the lungs when compared to an intratracheal delivery route [37]. Additionally, phages need to penetrate the lung epithelial barrier [38] in order to access the alveolar macrophages that are parasitized by *Mtb*, thus further increasing the difficulty of efficient intravenous phage administration.

To test this hypothesis, we compared phage copy number in spleens and in lungs, and the results indicated there are more phage copies in spleen than that in lungs. It is worth noting that our original dose contained 10^11^ phages per dose, while we only detected thousands of copies at the termination time point. This discrepancy may be due to the pharmacodynamical changes of the phages, as the phage’s half-life might be as low as 2.2 h and was undetectable after 36 h of injection [39, 40]. Even though we need to further investigate the pharmacokinetics of the phage DS6A, our qPCR result offered a reasonable explanation of why the intravenous administration of phage is more efficient in spleens than in lungs. Nevertheless, our results suggest that optimization of administration will be helpful to improve lung bacterial clearance (e.g., via intratracheal route or nebulizer delivery, etc.) as tuberculosis, in majority of cases, are infections of the lungs.

Phage resistance has always been a concern as it may dampen the therapeutic efficacy of phage therapy. In our study, it seems not to be a case for phage DS6A within the treatment period (Fig. 4f). However, in the clinical settings, the treatment time may be as long as several months, and *Mtb*-resistant strains may still appear. Therefore, screening more effective bacteriophage strains to form a phage cocktail would be essential to overcome phage resistance by *Mtb*.

Moreover, our study showed that significant IgM but weaker IgA and IgG responses were induced after 20 days of phage treatment (Fig. 5). This is because the short-term administration of phage may lead to IgM as dominant antibodies, and with the prolonged administration, the IgG response will become dominant. Given that phage treatment still improved disease progression in a human clinical case of *M. chelonae* despite a significant anti-phage antibody response [18], it may be that are not a great concern. Indeed, in our study, the antibody responses did not adversely affect the efficacy of phage therapy to eradicate *Mtb* in the spleen. This may be because our treatment regimen was not deployed long enough to enable to the development of sufficient quantity and quality of antibody responses to counteract the high dose of phages we administered. However, in clinical settings where a long treatment regimen might be needed, a high dose might be warranted to reduce the effects of antibody responses to the phages.

Taken together, our study showed that bacteriophage DS6A could effectively eliminate *Mtb* in agar plates, in liquid culture, in infected macrophages, as well as in humanized mouse model. Therefore, our study demonstrated the feasibility of developing phage therapy as an effective therapeutic against *Mtb* infection.

## Materials and Methods

### Regulatory and Ethical statement

Healthy donors were recruited for the collection of blood samples from the employees and students at the University of Texas Health Science Center at Tyler (UTHSCT) following the protocol approved by the Institutional Review Board (IRB) ethics committee of the UTHSCT. All animal procedures were approved by the UTHSCT Institutional Animal Care and Use Committee (IACUC).

### Bacterial strains, phage strains, media, and culture conditions

Bacteriophage DS6A was purchased from the ATCC (Cat# 25618-B2), and phages D29 and Chah were prepared at Queens College, The City University of New York (Which were originally obtained from Dr. Graham Hatfull’s lab at the University of Pittsburgh). *M. smegmatis* mc2 155 was purchased from the ATCC (Cat# 700084). *Mtb* H37Rv and *Mtb* H37Rv-GFP were prepared in our laboratories at UTHSCT. Except for DS6A, all mycobacteriophages were amplified in *M. smegmatis*; DS6A was amplified in *Mtb* H37Rv in the BSL-3 Laboratory at UTHSCT.

### Amplification and quantification of bacteriophages

Bacteriophages, except DS6A, were amplified in *M. smegmatis. M. smegmatis* was streaked on 7H10 agar plate supplemented with 10% Middlebrook Oleic Albumin Dextrose Catalase (OADC), 1mM CaCl_2_, 10 μg/mL of carbenicillin, and 100 μg/mL of cycloheximide and cultured at 37°C for about 3 - 4 days for single colony growth. A single colony of *M. smegmatis* was cultured in 7H9 media supplemented with 10% Middlebrook Albumin Dextrose Catalase enrichment supplement (ADC), 1mM CaCl_2_, 10 μg/mL of carbenicillin, 100 μg/mL of cycloheximide, and 0.25% Tween-80 at 37°C for 3- 4 days until the OD at 600 nm was over 2.0. A hundred mLs of *M. smegmatis* from the above culture was inoculated into 5 mL of 7H9 media supplemented with 10% ADC, 1mM CaCl2, 10 μg/mL of carbenicillin, 100 μg/mL of cycloheximide without Tween-80, and incubated overnight with shaking at 200 rpm at 37°C. Next day, 10^8^ pfu of bacteriophages with 100 μL of *M. smegmatis* were mixed and incubated at 37°C for 10 min, then the phage-bacilli mixture was added to 7mL of top agar at 60°C, and immediately applied onto 150-mm 7H10 agar plates. The phage-bacilli agar plates were incubated at 37°C overnight, then the bacilli were collected in 10 mL of phage buffer (10 mM Tris, pH 7.5, 10 mM MgSO4, 0.4% NaCl) and stored at 4°C overnight. After centrifugation, the supernatants were collected and filtered through a 0.22 μM PVDF filter, and the phages were stored at −80°C until use.

*Mtb* was cultured in 7H9 with 10% ADC following the standard *Mtb* culture procedures. *Mtb* single cell suspensions at 10^8^ cells per ml were infected with phage DS6A using an MOI of 1 for 10 min at 37 °C. The infection mixture was mixed with 7 mL of top agar (0.8% agarose in 7H9 with 10% ADC) and applied onto 150 mm 7H10 agar plate (supplemented with 10% OADC, 1 mL of CaCl_2_). The plates were incubated at 37°C for ∼8 days. After plaques formed, the phage DS6A was collected in phage buffer and filtered through a 0.22 μM PVDF filter.

Bacteriophages’ titers were determined by plaque assays by infection of H37Rv with phage DS6A or *M. smegmatis* with other phages, respectively, after serially diluted the phages. Alternatively, quantitative PCR was also used to quantify the viral genomes.

### Testing *Mtb*-killing ability on 7H10 agar plates

To test whether the bacteriophages can kill *Mtb* on the solid agar plate culture, we performed the plaque assays. Briefly, 10^4^, 10^3^, 10^2^ pfu of D29, Chah (for both the titers based on *M. smegmatis*), and DS6A (the titer based on *Mtb*) were mixed with 1 x 10^5^ *Mtb* bacilli and incubated at 37°C for 1 h with shaking. The *Mtb*-phage mixture was then mixed with 500 µL of 0.8% top agar in 7H9 with 10% ADC and applied to 12-well 7H10 agar plates. The plates were incubated at 37°C for ∼7-8 days until the plaques developed.

### Testing *Mtb*-killing ability in 7H9 liquid culture

The *Mtb* killing activities of the bacteriophages were determined by infecting 1 x 10^5^ H37Rv in 1 mL 7H9 media with various bacteriophages at an MOI of 1 for 10 min, then inoculated into 20 mL 7H9 media supplemented with 10% ADC and incubated at 37°C for 9 days. At days 3, 6, and 9, 10 μL of 10-fold serial dilutions of the culture were plated on 7H10 agar plates, and CFU were counted after cultured at 37°C for 2-3 weeks.

### Flow cytometry

Cells from one well were collected and stained for macrophage markers. The antibodies used were APC-conjugated CD11b (cat# 301350), FITC-conjugated CD14 (cat# 325604), BV711-conjugated CD16 (cat# 302044). The isotype controls included APC-mouse IgG1 (cat# 400142), FITC-mouse IgG1 (cat# 400108), BV711-mouse IgG1 (cat# 400168). All antibodies were purchased from BioLegend. Briefly, the cells were washed with PBS and incubated with the antibodies at room temperature for 15 min, avoiding light. The cells were then washed and resuspended with a buffer of 0.5% BSA/PBS + 3mM EDTA prior to analysis with Attune NXT (ThermoFisher). Compensation was performed using UltraComp eBeads Plus (01-3333-42, Invitrogen). The data were then analyzed with FlowJo version 10.6.1, and the graphs were created with GraphPad Prism 8.

### Testing the *Mtb*-killing ability of the bacteriophages in macrophages

Blood was drawn from five healthy donors and PBMCs were isolated using Ficoll-Paque Plus (17-1440-02, GE Healthcare, Danderyd, Sweden) gradient centrifugation. Then, CD14+ cells were isolated from the PBMCs using human CD14+ micro beads conjugated with anti-human CD14 mAbs (130-118-906, Miltenyi Biotec, Bergisch Gladbach, Germany) LC columns (130-042-401, Miltenyi Biotec, Bergisch Gladbach, Germany) following the manufacturing instructions. The purity of the cells was checked with flow cytometry.

Half a million cells were seeded in each well of 24-well plates containing 1.5 mL of RPMI-1640 supplemented with 10% FBS and 1% penicillin + streptomycin. The cells were then incubated in the presence of 50 ng/mL of GM-CSF (02532, Stem Cells, Vancouver, Canada) and 50 ng/mL of M-CSF (216-MC, R&D Systems, Minneapolis, USA) at 37°C and 5% CO_2_ for differentiation. Fresh cytokines were replenished on day 3 once. On day 6, some cells were collected and stained for macrophage markers followed by flow cytometry analysis. The rest of the cells were washed with PBS and rested overnight in fresh medium without serum and antibiotics. The cells were then infected with H37Rv at an MOI of 1. Four hours post infection, the cells were washed with warm PBS thrice to remove extracellular bacilli. The cells were then incubated in fresh RPMI-1640 with 10% FBS with no antibiotics as controls and with either DS6A or D29 control phage at 37°C. The growth of H37Rv in macrophages were followed for different time points after infection by plating serially diluted cellular lysates on 7H10 agar plates as follows: After removing the culture medium, the cells were treated with ddH_2_O for 10 minutes followed by 0.25% SDS in HBSS for 10 min. At the end, 20% BSA in PBS was added to stop the lysis. The content of each well was collected into a tube and sonicated three times with 33% amplitude for 10 seconds. After making dilutions, 10 µL of cellular lysate was plated on 7H10 agar plates in duplicates. The plates were incubated at 37°C for 2-3 weeks and the bacterial colonies were counted for CFU determination.

### Generation of humanized mice

Humanized NSG-SGM3 mice (NSG mice transgenically expressing three human cytokine/chemokine genes IL-3, GM-CSF, and KITLG, RRID: IMSR_JAX:013062) were obtained from Jackson Laboratories and bred in UTHSCT vivarium. To increase scientific rigor, the experiments have been done in a blinded manner. To exclude sex bias, each experiment was considered to contain an equal number of male and female mice. For humanization of NSG-SGM3 mice, the mice were irradiated with a dose of 100 cGy per mouse and injected intravenously with 0.2 million of HSCs (purchased from Stemcell Technologies Inc.). After 8–10 weeks, the human cell reconstitution was confirmed by flow cytometry analysis of human immune cells in the mice by staining the PBMCs with fluorescent human antibodies specific for CD45, CD3, CD4, CD8, CD14, CD19, and CD56.

### *Mtb* infection and phage treatment of humanized mice and other animal experiments

Humanized mice were infected with a low dose of aerosolized H37Rv in a Madison chamber as previously described [41] by properly diluted H37Rv stocks in 10 mL normal saline to deposit about 100 CFUs per mouse lung as confirmed by the determination of Lung CFUs 1 day post infection. All animal experiments were done in a blinded manner to increase scientific rigor. Wherever possible, each experiment contains equal numbers of male and female mice to exclude sex bias.

*Mtb*-infected mice were treated with 1 x 10^11^ pfu phage DS6A per mouse or equal volumes of PBS as a control by intravenous administration every other day for a total of 8 treatments. The mice’s body weight was measured every other day during the treatment. Three days after the last treatment, the mice were subjected to pulmonary function testing and CT scan as described previously [42]. Briefly, mice were anesthetized with a ketamine/xylazine mixture, and the anesthetized mice were cannulated with a sterile, 20-gauge intravenous cannula through the vocal cords into the trachea. The snapshot perturbation method was used to evaluate lung compliance by collecting the measurements of elastance, compliance, and total lung resistance for each mouse using the FlexiVent system (SCIREQ, Tempe, AZ). The flexiVent was set to a tidal volume of 30 mL/kg at a frequency of 150 breaths/min against 2–3 cm H_2_O positive end-expiratory pressure, according to manufacturer’s instructions.

After lung function testing, the mice were immediately subjected to CT scans for the measurements of lung volume. The Explore Locus Micro-CT Scanner (General Electric, GE Healthcare, Wauwatosa, WI) was used for CT imaging. CT scans were performed during full inspiration and at a resolution of 93 μm. Lung volumes were calculated from lung renditions collected at full inspiration. Microview software 2.2 (http://microview.sourceforge.net) was used to analyze lung volumes and render three-dimensional images. The mice were maintained under anesthesia using isofluorane throughout the pulmonary function testing.

At the end of testing, the mice were euthanized, and the lungs and spleens were collected and homogenized. The serially diluted lung and spleen homogenates were plated on 7H10 agar plates with OADC and incubated at 37°C for 2-3 weeks for the determination of the organ CFUs.

### ELISA for detection of phage-specific antibodies

Phage DS6A were coated in 96-well Nunc-Immuno plates (Thermofisher) with a concentration of 5×10^9^ pfu/mL in coating buffer (Na2CO3, pH8.0) at 4°C overnight. After washing with PBST (0.1% Tween-20 in PBS) four times, the plate was blocked with PBSM (5% skimmed milk in PBS) and followed by applying 10-fold serially diluted humanized mouse sera and incubated for 2 h at room temperature, then washed with PBST for 5 times. 100 mL of HRP-labeled secondary antibodies were applied to the plates and incubated at room temperature for 1 h. The secondary antibodies included: (1) goat anti-human IgG Fc (HRP) pre-adsorbed (catalog no. ab98624; Abcam); (2) goat anti-human IgA alpha chain (HRP) pre-adsorbed (catalog no. ab98558; Abcam); and (3) goat anti-human IgM mu chain (HRP) pre-adsorbed (catalog no. ab98549; Abcam). After washing away the unbound secondary antibodies, TMB substrate was added and the OD values at 450 nm were recorded using a BioTek Synergy 2 plate reader.

### Multiplex assay for cytokine profiling

Lung and spleen tissue samples from both experimental groups were homogenized in PBS to a final volume of 1.5mL, using a 70 µm cell strainer (MTC Bio, Sayreville, NJ). The homogenate was passed through a 0.2 µm filter (Sigma-Aldrich, St. Louis, MO), and the flowthrough was used to evaluate the cytokine profile in these tissues. The samples were analyzed in duplicates using the Bio-Plex Pro™ Human Cytokine panel (Bio-Rad, Hercules, CA), according to the manufacturer’s instructions. Briefly, 50 µL of filtered tissue homogenate were dispensed in a 96-well plate containing magnetic beads conjugated with antibodies for the detection of 27 different cytokines. After further incubation with detection antibodies and streptavidin-PE, the samples were analyzed in the Bio-Plex MAGPIX multiplex reader (Bio-Rad Laboratories Inc., CA). Fluorescence values were converted into cytokine concentrations (expressed as pg/mL), using a regression curve based on the values obtained from a set of standard dilutions.

The cytokines reported by the Bio-Plex Pro™ Human Cytokine panel were: Basic FGF, Eotaxin, G-CSF, GM-CSF, IFN-γ, IL-1β, IL-1Ra, IL-2, IL-4, IL-5, IL-6, IL-7, IL-8, IL-9, IL-10, IL-12, IL-13, IL-15, IL-17, IP-10, MCP-1, MIP-1α, MIP-1β, PDGF-BB, RANTES, TNF-α and VEGF.

### Statistical analysis

Each treatment was triplicated (PCR) or duplicated (all other experiments), and the experiments were repeated at least once to ensure reproducibility. Power analysis was done to determine the sample size to ensure biological significance. The data were analyzed using GraphPad Prism software. Unpaired student T-tests were used to analyze the differences between treated and control groups. All statistical data are represented as mean ± SEM. Statistical significance was defined as *P≤0.05, **P≤0.01, and ***P≤0.001.

## Data availability

All data supporting the findings of this study are available in the manuscript. If there are any special requests or questions for the data, please contact the corresponding author (G.Y.).

## Acknowledgments

We thank Dr. Amy Tvinnereim for helping perform the following experiments: *Mtb* infection of the humanized mice and the pulmonary function test.

## Contributions

Guohua Yi: Conceived and guided the study, designed and performed experiments, analyzed data, edited figures, and wrote and finalized the manuscript.

Fan Yang: Performed the experiments and analyzed data.

Alireza Labani-Motlagh: Performed the experiments and analyzed data.

Josimar Dornelas Moreira: Performed the experiments.

Danish Ansari: Performed the experiments.

Jose Alejandro Bohorquez Garzon: Performed the experiments.

Sahil Patel: Performed the experiments.

Fabrizio Spagnolo: Performed the experiments and edited the manuscript.

Jon Florence: Performed the experiments.

Abhinav Vankayalapati: Performed the experiments.

Ramakrishna Vankayalapati: Participated in experiment discussions, provided suggestions and edited the manuscript.

John J. Dennehy: Designed experiments, participated in experiment discussions, provided phage strains, and edited the manuscript.

Buka Samten: Designed experiments, guided experiments, performed experiments, participated in experiment discussions, and edited the manuscript.

