## Supplementary materials for "Bacteriophage therapy for the treatment of *Mycobacterium tuberculosis* infections in humanized mice"

**Supplementary Figures**

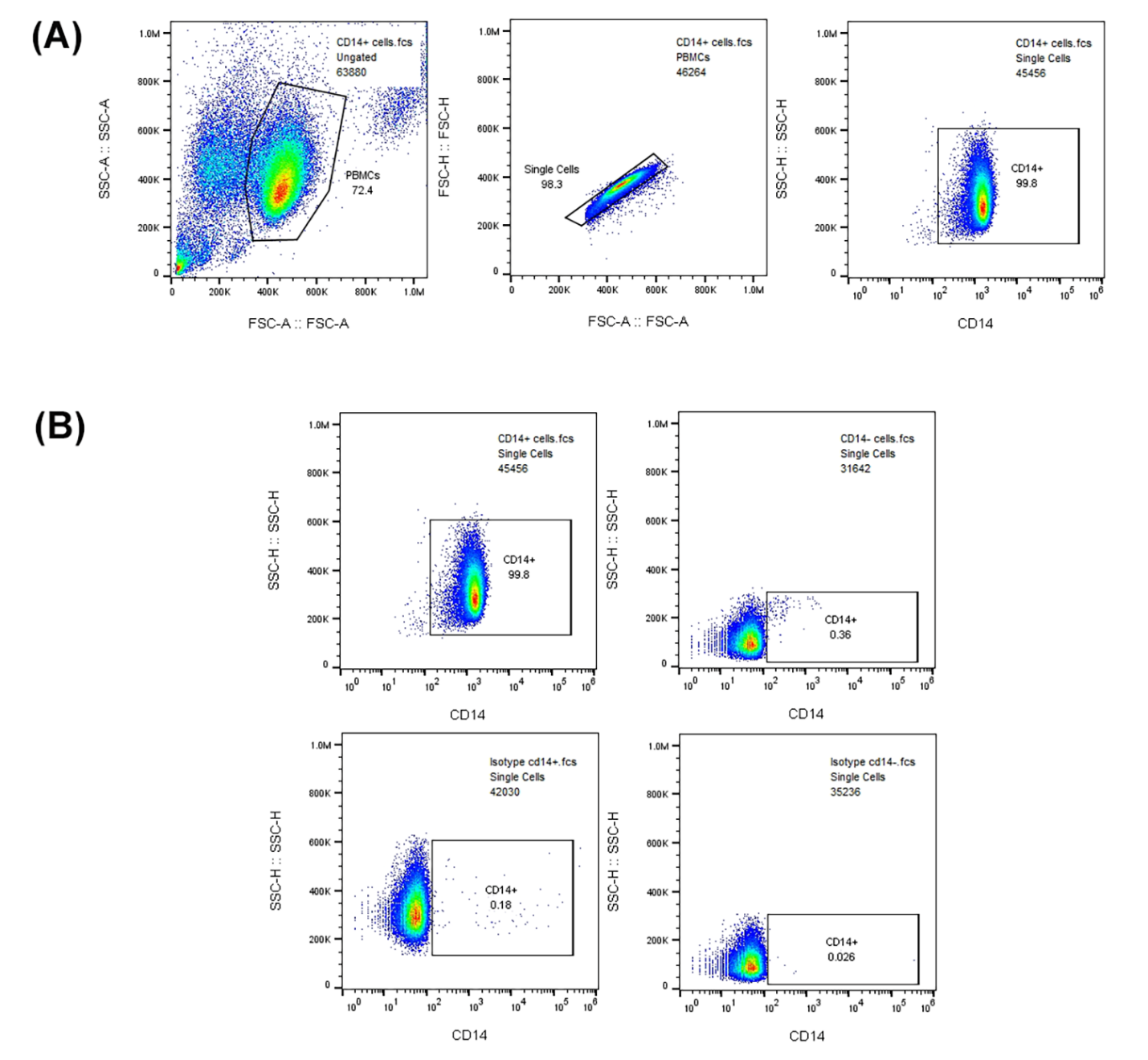

**Supplementary figure 1:** The purity of microbeads isolated CD14+ cells. (A) Gating strategy. (B) The purity of CD14+ cells. The top panel shows the percentages of CD14+ cells (left) and CD14- cells (right). The bottom panel shows the isotype controls of CD14+ and CD14- cells.

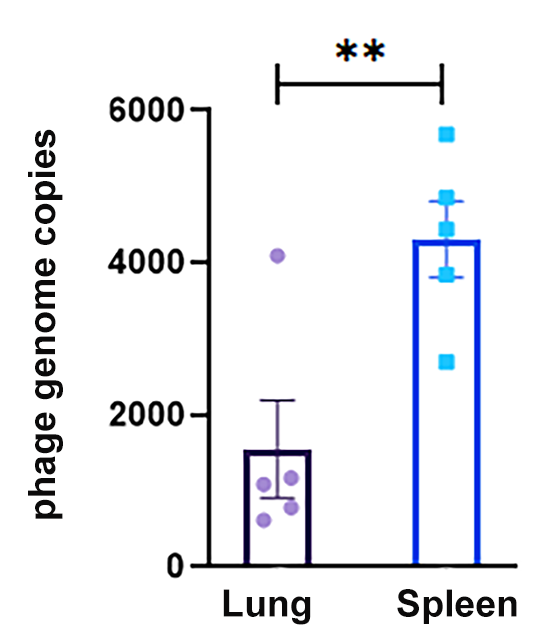

**Supplementary figure 2:** Phage genomic copy numbers in the lung and spleen homogenates of the phage-treated humanized mice. Five phage-treated mouse lung and spleen homogenates were used to isolate phage genomic DNA, and quantitative PCR was performed to determine the phage copy numbers in the tissue homogenates. Each dot represents the mean of triplicates, and paired T test was used to analyze the differences between the lung and spleen homogenates. Statistical significance was defined as *P≤0.05, **P≤0.01, and ***P≤0.001.

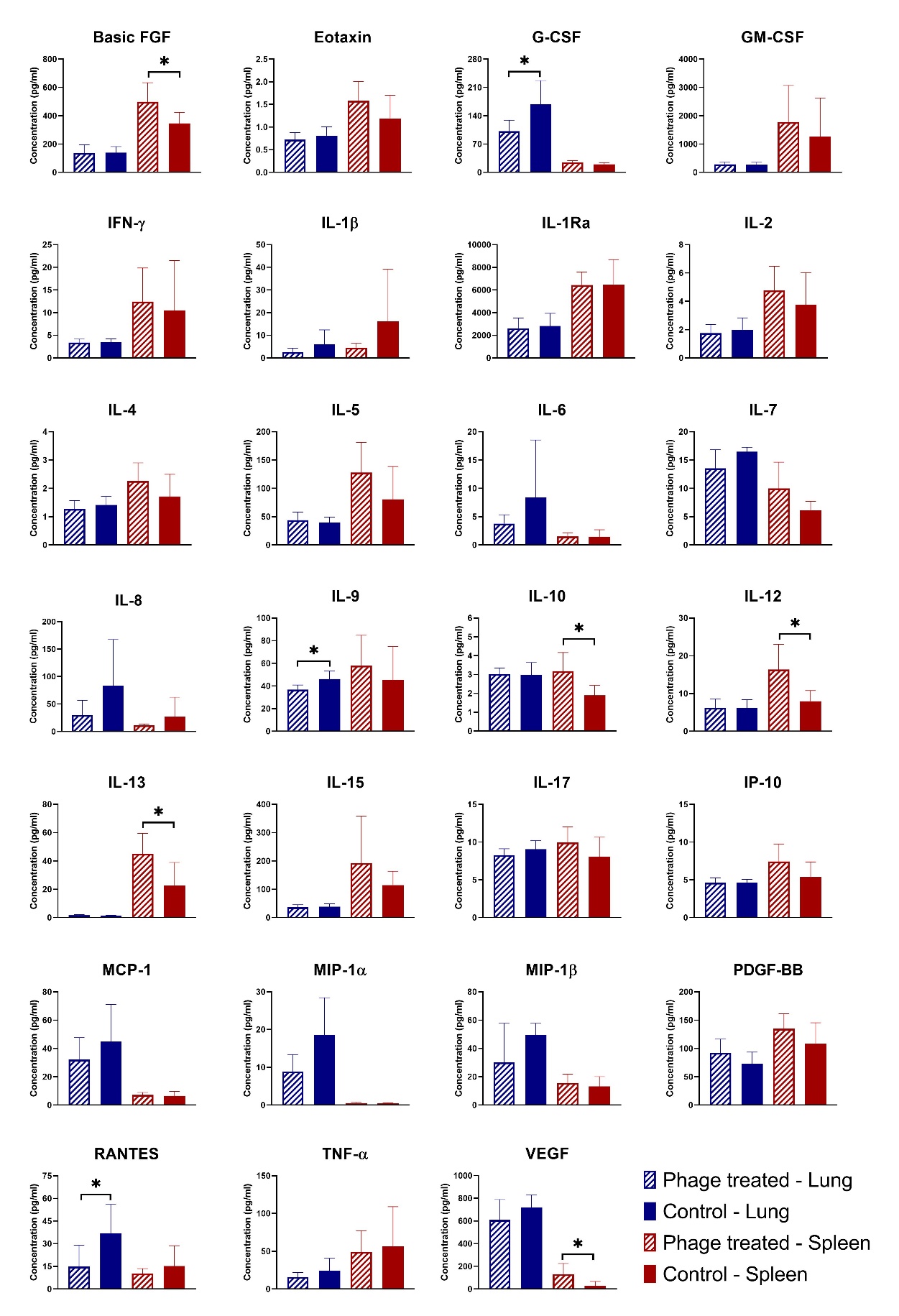

**Supplementary figure 3:** Cytokine profiles of the experimental mouse spleen and lung homogenates. Twenty-seven cytokines and chemokines of human lymphocytes and myeloid cells in the lung and spleen homogenates of the humanized mice were determined by multiplex assay. Five mice from control group (*Mtb*-infected) and phage treated group were used for cytokine profiling. Unpaired T test was used to analyze the differences between control and phage-treated group. Statistical significance was defined as *P≤0.05, **P≤0.01, and ***P≤0.001.
